# Unexpected ribosome turnover during prolonged translation inhibition

**DOI:** 10.64898/2026.05.06.723260

**Authors:** Paul J. Russell, Caleb A. Clark, Mia Ashreim, Michael G. Kearse

**Affiliations:** Department of Biological Chemistry and Pharmacology, Center for RNA Biology, The Ohio State University, Columbus, OH 43210

## Abstract

Eukaryotes use several distinct quality control pathways to resolve aberrant ribosomes and mRNAs. For example, the no-go decay mRNA pathway is stimulated after ribosome collisions caused by stalled ribosomes translating damaged or truncated mRNAs. Separate decay pathways for non-functional 40S and 60S subunits containing rRNA mutations affecting decoding and peptidyl transferase activity, respectively, have also been elucidated. To our knowledge, whether eukaryotes have evolved a quality control pathway to sense and process globally stalled ribosomes is unclear; however, such a pathway would be advantageous to eukaryotes during exposure to natural elongation inhibitors such as ricin and diphtheria toxin. Here, we test how prolonged robust inhibition of elongation using a high dose of cycloheximide (CHX) affects ribosome turnover. Despite no decrease in cell viability and that mammalian ribosomes have been classically characterized of having a half-life of 3-5 days, a single 24 hr high dose of CHX resulted in drastically shortened half-lives (<24 hr) of 28S and 18S rRNA in A549 cells. A ∼2-fold reduction in nearly all ribosome species was observed by polysome analysis in HeLa and A549 cells after prolonged CHX treatment. Depletion of ribosomes was also evident when assessing ribosomal proteins from both the 40S and 60S subunits by Western blot. Literature supports that ribosomes can be degraded by autophagy and the ubiquitin (Ub)-proteasome system. Upon testing inhibitors of both pathways, only proteasome inhibitors (*i.e.*, MG132 and bortezomib) rescued both rRNA and ribosomal protein levels. Proteasome inhibitors also rescued ribosome levels in polysome profiling experiments. Remarkably, rRNA levels were not rescued during CHX treatment when co-treated with the Ub activating enzyme E1 inhibitor, TAK243. Polysome analysis also showed that the high prolonged dose of CHX did not cause robust accumulation of collided ribosomes compared to control treatments. Proteasome-dependent turnover of rRNA was also observed with high doses of other elongation inhibitors, namely anisomycin, homoharringtonine, and lactimidomycin. The recognition capabilities of the pathway were further expanded as we observed that 80S ribosomes not trapped on the mRNA were also targeted for degradation by the proteasome. Together, our findings define the framework of a regulatory pathway in mammalian cells that degrades both ribosomal subunits in response to prolonged periods of robust elongation inhibition. We refer to this novel pathway as the Global Ribosome Instability Pathway (GRIP).

## INTRODUCTION

Ribosomes are classically thought of as the protein factories of the cell. Indeed, canonical eukaryotic translation elongation is the process where ribosomes in coordination with aminoacylated-tRNAs decode the sense codons of the open reading frame (ORF) to produce a gene specific polypeptide. An emerging function of ribosomes is their ability to act as cellular sensors. A well-studied example of this is the ribosome-associated quality control (RQC) pathway. This pathway has evolved to remove aberrant mRNAs from the cell that if otherwise translated would produce nonfunctional and potentially gain-of-function toxic proteins. During the process of translation elongation, if the ribosome encounters obstructions, it can lead to ribosome stalling. When a leading ribosome stalls on the mRNA and is left unresolved, the trailing ribosome will impact it and result in a ribosome collision. The collision event stimulates activation of the RQC pathway by creating a disome interface that is recognized by an E3 ubiquitin ligase, ZNF598 (1,2). Ubiquitylation of ribosomal proteins (RP) on the 40S ribosomal subunit RPS10 and RPS20 results in downstream recruitment of factors to the ribosome to degrade the nascent polypeptide and the mRNA (3). In the case of the mRNA, it is believed that following ubiquitylation of RPS10 and RPS20, Cue2 in yeast (the mammalian ortholog is unknown but speculated to be N4BP2) is recruited to the ribosome (4). Endonucleolytic cleavage of the mRNA produces free 5’ and 3’ ends that can then be degraded by Xrn1 and the exosome complex, respectively (5). Following degradation of the mRNA, the nascent polypeptide is subsequently targeted for turnover. This occurs after the 80S ribosome is split into its separate subunits (6,7). The obstructed 60S ribosomal subunit is recognized by NEMF which aids in the recruitment of another E3 ubiquitin ligase, listerin, to the nascent polypeptide (8–10). Listerin ubiquitylates the nascent polypeptide allowing for recognition and degradation by the proteasome (11–16). Importantly, the canonical RQC pathway promotes ribosome recycling and does not lead to apparent ribosome degradation.

In contrast to the RQC pathway, under stress conditions that impact global translation, the ribotoxic stress response (RSR) pathway is activated. It has been shown that the RSR pathway is stimulated following amino acid starvation or treatment with UV irradiation, ribotoxins, and elongation inhibitors (17–20). Following exposure to these stress conditions, ribosome collisions predominately occur (17,20) although there is some evidence that stalled monosomes can also activate the RSR pathway (18,19). Instead of ZNF598 recruitment to the collided disomes, ZAKα, which generally associates with elongating ribosomes (17,19), senses the collision and becomes activated through autophosphorylation (19). Consequently, phosphorylated ZAKα dissociates from the ribosome (20) and leads to downstream phosphorylation and activation of stress-activated protein kinases (SAPKs), p38 and c-Jun N-terminal kinase (JNK), which mediate cell cycle arrest and cell apoptosis, respectively (17–20). Additionally, ZAKα activation during periods of low and intermediate stress triggers activation of an Integrated Stress Response (ISR) kinase, GCN2 (17,20). This results in phosphorylation of the alpha subunit of eukaryotic initiation factor (eIF) 2 and subsequent downregulation of translation. Repression of global translation by eIF2α phosphorylation prevents additional recruitment of ribosomes to mRNA and limits further ribosome collisions. Recent work has shown that the severity of ribosome collisions determines cell fate (20). Under conditions when ribosomes collisions are limited, p38 and GCN2 activation predominate as measures to alleviate the stress and return the cell to normal homeostasis. Alternatively, when ribosome collisions are abundant and persistent, robust activation of JNK occurs and downstream signaling leads to cell apoptosis.

One caveat to these pathways is that little evidence exists for their functional relevance after prolonged periods of translational stress as most quality control pathways resolve ribosome collisions on short time scales. One translation elongation inhibitor commonly used to block protein synthesis or to measure translation efficiency of mRNAs is cycloheximide (CHX). Inhibition of translation results from CHX binding to the E site of the 60S ribosomal subunit, which consequently prevents translocation of the ribosome along the mRNA (21). It has previously been shown, when HeLa cells were treated with high dosages of CHX for 24 hrs, that nearly half of all ribosomal species were degraded (22). It was determined that the reduction of ribosomes was not due to loss of cell viability or changes to cellular metabolism. This has also been corroborated by recent work that specifically showed that HeLa cells in interphase treated with similar high concentrations of CHX survived between 1-2 days in culture (23). Interestingly, HeLa cells in the mitotic phase treated with CHX only survived for approximately 4 hrs (23). Considering that these cells remain viable and do not undergo bulk cell apoptosis under these conditions, it stands to reason that globally stalled ribosomes would be degraded through an unknown quality control pathway. Here we show that, following prolonged periods of translation elongation inhibition, almost half of mature ribosomes in A549 cells are degraded. We further demonstrate that the turnover of these ribosomes is through a proteasome-dependent mechanism that most likely does not require site-specific ubiquitylation of RPs. Additionally, we show that the pathway indiscriminately targets ribosomes as ribosomes trapped in different states on the mRNA and those dissociated from the mRNA are all subject to proteasome-dependent turnover. We also provide evidence that similar levels of rRNA degradation are observed following treatment with lysosome permeabilization agents; however, treatment with CHX does not result in lysosome membrane permeabilization. Moving forward, we aim to further mechanistically define the mature ribosome turnover pathway and identify the factors required for RP and rRNA degradation. We refer to this novel pathway as the Global Ribosome Instability Pathway (GRIP).

## MATERIALS AND METHODS

### Cell culture and drug treatments

A549 cells were obtained from ATCC (catalog # CCL-185) and maintained in high glucose DMEM (Thermo Fisher #11995065) supplemented with 10% heat-inactivated Fetal Bovine Serum (Cytiva # SH30396.03), MEM Non-Essential Amino Acids (Thermo Fisher #11140-050), 1% Penicillin-Streptomycin (Thermo Fisher # 15140122) at 37°C and 5% CO_2_ in standard tissue culture treated plates.

The following inhibitors were used in this study: CHX (Sigma #C1988; 100 mg/mL stock in DMSO), PURO (Sigma #P9620; 10 mg/mL stock in water), ANS (Sigma #A5862; 10 mg/mL stock in DMSO), HHT (Sigma #SML1091; 10 mg/mL stock in DMSO), LTM (Millipore #506291; 5 mM stock in DMSO), CX-5461 (Sigma # 5092650001; 1 mM stock in DMSO), TAK243 (Selleckchem #S8341; 10 mM stock in DMSO), MG132 (Sigma #M7449; 10 mM stock in DMSO), Bortezomib (Selleckchem #PS341; 10 mM stock in DMSO), SAR405 (Cayman Chemical #16979; 10 mM stock in DMSO), Baf A1(abcam #ab120497; 300 µM stock in DMSO), PLX-4720 (Selleckchem #S1152, 100 mM stock in DMSO), L-Leucyl-L-Leucine methyl ester (hydrochloride) (Cayman Chemical #16008; 1 M stock in DMSO), and Z-VAD-FMK (Selleckchem #S7023; 50 mM stock in DMSO).

### RNA interference

4 μL siRNA (neg # 1; Thermo Fisher # 4390843; neg # 2; Thermo Fisher # 4390847; ZAK # 1; Thermo Fisher # 4427038, ID: s28651, or ZAK # 3; Thermo Fisher # 4427038, ID s28653) was added to 6-well plates. 5.25 μL lipofectamine RNAiMAX (Invitrogen # 13778150) and 1.5 mL Opti-MEM (Gibco # 31985-020). was then added to the wells. The mixture was incubated at room temperature for 30 min. 1.5mL of A549 cells was added to the wells and plates were incubated overnight at 37°C. Media was aspirated the next day and 3 mL of media was added back to the wells. After an additional two days (72 hrs post transfection) when cells were ∼50% confluent, wells were washed with 1 mL cold PBS, and 600 μL RIPA buffer with protease inhibitor EDTA-free (Thermo Fisher # A32955) and phosphatase inhibitor EDTA free (Thermo Fisher # A32957) was added for 10 min with gentle shaking at 4°C.

### *In vitro* transcription

nanoLuciferase (nLuc) mRNA was synthesized using linearized plasmids as templates for run off transcription with T7 RNA polymerase as previously described (24) with the single exception that XbaI was used to linearize all plasmids. mRNA was transcribed at 30°C for 2 hrs using the HiScribe T7 High Yield RNA Synthesis Kit (NEB # E2040S). mRNA was purified using the Zymo RNA Clean and Concentrator-25 (Zymo Research # R1018), eluted in RNase-free water, aliquoted in single use volumes and stored at −80°C.

### RNA Denaturing Gels

A549 cells were seeded into 6-well plates with 3 mL of media. After 72 hrs, cells (∼50% confluency) were treated with inhibitors. 24 hrs post-treatment, the media was aspirated, and total RNA was extracted from each well by adding 1 mL TRIzol (Thermo Fisher # 15596018) and following the manufacturer’s protocol. The resulting RNA pellet was resuspended in 20 μL nuclease-free water. Equal volume of total RNA (based on 250 ng RNA for 0 hr control) containing 100 ng of spike-in nLuc mRNA as a loading control was separated on a 1% denaturing formaldehyde agarose gel in 1X MOPS buffer. Gels were imaged on the BioRad GelDoc Go Imaging System using the Ethidium Bromide setting. 28S and 18S rRNA levels were normalized to nLuc.

### 4SU Labeling

A549 cells were seeded in 10 cm plates with 10 mL of media. After 72 hrs, cells (∼50% confluency) were pre-treated with either vehicle (DMSO), 100 μg/mL CHX or 2.5 μM CX-5461 for 1 hr at 37°C. Following pretreatment, cells were treated with vehicle (DMSO) alone or co-treated with 10 μM 4SU and either vehicle (DMSO), 100 μg/mL CHX or 2.5 μM CX-5461 for 16 hrs at 37°C. Total RNA was extracted from each plate by adding 3 mL TRIzol (Thermo Fisher # 15596018) and following the manufacturer’s protocol. The resulting RNA pellet was resuspended in 20 μL nuclease-free water. Total RNA was biotinylated using 1 mg/mL EZ-Link HPDP-Biotin for 2 hrs at 25°C in the dark. RNAs were then purified using the Zymo RNA Clean and Concentrator-5 (Zymo Research # R1015), eluted in 20 μL RNase-free water, and stored at − 80°C. 4 μg of total RNA was separated on a 1% denaturing formaldehyde agarose gel. RNA was transferred to BrightStar-Plus Positively Charged Nylon Membrane (Thermo Fisher #AM10104) using a previously described method (25). Transfer apparatus was then disassembled, and the nylon membrane was UV crosslinked using the Stratagene Stratalinker UV 1800 crosslinker with 120,000 µJ/cm2 (Auto Crosslink function). Total RNA was subsequently detected by staining with 0.25% (w/v) methylene blue (in 0.4 M sodium acetate and 0.4 M acetic acid) for 15 min at room temperature with orbital shaking, washed with distilled water, and then imaged on the BioRad GelDoc Go Imaging System using the Coomassie Blue setting. The same membrane was then washed with more distilled water to remove methylene blue stain. Afterwards, the membrane was washed with 2XSSC, 0.1% SDS for 15 min at room temperature with orbital shaking. Chemiluminescence was performed using Chemiluminescent Nucleic Acid Detection Module (Thermo # 89880) and following the manufacturer’s protocol. Images were taken using an Azure Sapphire Biomolecular Imager.

### rRNA decay measurements

A549 cells were seeded in 6-well plates with 3 mL of media. After 72 hrs, cells (∼50% confluency) were treated with either vehicle (DMSO), 100 μg/mL CHX or 2.5 μM CX-5461. Cells were collected at 6 hr timepoints for 24 hrs. Total RNA was extracted from each well by adding 1 mL TRIzol (Thermo Fisher # 15596018) and following the manufacturer’s protocol. The resulting RNA pellet was resuspended in 30 μL nuclease-free water. Equal volume of total RNA (based on 250 ng RNA for 0 hr control) containing 100 ng of spike-in nLuc mRNA as a loading control was separated on a 1% denaturing formaldehyde agarose gel in 1X MOPS buffer. Gel was imaged on the BioRad GelDoc Go Imaging System using the Ethidium Bromide setting. 28S and 18S rRNA levels were normalized to nLuc and half-lives were calculated by simple linear regression in GraphPad Prism 10.3.1.

### Western blot analysis

A549 cells were seeded into 12-well plates with 1.5 mL of media. After 24 hrs, cells (∼50% confluency) were treated with inhibitors. 24 hrs post-treatment, the media was aspirated, cells were gently washed with 1 mL ice-cold PBS and then lysed with 300 µL ice-cold RIPA buffer containing protease inhibitors. Plates were incubated for 10 min at 4°C with gentle rocking. Samples were prepared by adding 4X reducing Laemmli sample buffer (Bio-Rad # 1610737), heating for 15 min at 70°C, and syringing with 28G needles. 20-30 µL was then separated by Tris-Glycine SDS-PAGE (Invitrogen # XP04200BOX or XP04205BOX) and transferred on to 0.2 µm PVDF membrane (Thermo # 88520). Membranes were then blocked with 5% (w/v) non-fat dry milk in TBST (1X Tris-buffered saline with 0.1% (v/v) Tween 20) for 30 min at room temperature before overnight incubation with primary antibodies in TBST or in 5% (w/v) non-fat dry milk in TBST at 4°C with gentle rotation. Membranes were then washed three times with TBST for 10 min before incubation with an HRP-conjugated secondary antibody in TBST for 1 hr at room temperature. Membranes were then washed three times with TBST for 10 min. Chemiluminescence was performed using SuperSignal West Pico PLUS (Thermo # 34577) and imaged using an Azure Sapphire Biomolecular Imager.

Mouse anti-Actin (Cell Signaling # 3700S), mouse anti-alpha (α) Tubulin (Sigma # T9026), rabbit anti-Cyclin D1 (Cell Signaling #55506S; clone E3P5S), rabbit anti-p38 MAPK (Cell Signaling #9212S), rabbit anti-P-p38 MAPK (Cell Signaling #9211S, clone T180/Y182), rabbit anti-RPS6 (Cell Signaling # 2217; clone 5G10), rabbit anti-RPL7 (abcam #ab72550), rabbit anti-RPS3 (Bethyl #A303-840A), rabbit anti-RPL23A (Bethyl #A303-932A), rabbit anti-Ubiquitin (Cell Signaling #43124; clone E4I2J), and rabbit anti-ZAK (Bethyl #A301-993A) were used at 1:1000 in TBST with 0.02% (w/v) sodium azide. HRP-conjugated goat anti-rabbit IgG (H+L) (Thermo #31460) was used at 1:10,000 for anti-Cyclin D1, -p38 MAPK, -P-p38 MAPK, -Ubiquitin, and - ZAK and 1:20,000 for anti-RPS3, -RPS6, -RPL7, and -RPL23A. HRP-conjugated goat anti-mouse IgG (H+L) (Thermo #31430) was used at 1:10,000 for anti-Actin and -alpha (α) Tubulin.

### Sucrose gradient ultracentrifugation

Per replicate, A549 cells were seeded in two 15 cm tissue culture treated plates and allowed to grow for 72 hrs until ∼50% confluent. Cells were treated for 15 min with 100 μg/mL CHX or for 24 hrs with either 100 μg/mL CHX or 100 μg/mL CHX and 10 μM MG132. For collection, cells were gently washed with ice-cold PBS with 100 µg/mL cycloheximide or 100 µg/mL CHX and 1 µM MG132. After washing, additional ice-cold PBS with 100 µg/mL CHX or 100 µg/mL CHX and 1 µM MG132 was added to the plate. Cells were harvested using a cell lifter, pooled, pelleted at 500 rcf for 5 min at 4°C, snap froze in liquid nitrogen, and stored at -80°C. Cell pellets were thawed on ice and then lysed in 450 μL ice-cold Polysome Lysis Buffer (20 mM Tris-HCl, 140 mM KCl, 10 mM MgCl_2_, 1 mM DTT, 100 μg/mL CHX, 1% (v/v) Triton X-100; pH 7.4) for 10 min on ice. Cell debris was then cleared at 13,000 rcf at 4°C in a pre-chilled microcentrifuge. The supernatant was taken and 450 µL was aliquoted into two new pre-chilled microcentrifuge tubes. For RNase treated samples, 0.5 mM CaCl_2_ (final) followed by 150 U of S7 micrococcal nuclease (Roche #10107921001; stock at 30 U/µL in PBS) was added to cleared lysates, incubated at 25°C for 10 min, and then quenched by adding 1 mM EGTA (final). 400 µL was then layered on top of a linear 10-50% (w/v) buffered sucrose gradient (20 mM Tris-HCl, 140 mM KCl, 10 mM MgCl_2_, 1 mM DTT, 100 μg/mL CHX; pH 7.4) in a 14 mm × 89 mm thin-wall Ultra-Clear tube (Beckman # 344059) that was formed using a Biocomp Gradient Master. Gradients were centrifuged at 35K rpm for 120 min at 4°C in a SW-41Ti rotor (Beckman) with maximum acceleration and no brake using a Beckman Optima L-90 Ultracentrifuge. Gradients were subsequently fractionated using a Biocomp piston fractionator with a TRIAX flow cell (Biocomp) recording a continuous A_260_ nm trace.

### Denaturing PAGE

8% PAGE gels were made with SequaGel UreaGel 19:1 Denaturing Gel System (National Diagnostic #EC-833). Gels were poured between glass plates and allowed to polymerize for at least 1 hr. Gels were pre-run for 20 min at 100 volts (constant) with 1X TBE Running Buffer. Equal volume of total RNA (based on 1 μg RNA for 0 hr control) was loaded, and gels were run for 2 hrs at 100 volts (constant) with 1X TBE Running Buffer. Total RNA was stained with 1X SYBR Green II RNA Gel Stain (Thermo Fisher # S7568) diluted in milliQ water for 40 min in the dark. Stained gels were imaged on a Bio-Rad GelDoc Go Gel Imaging System using the SYBR Green setting.

### Fluorescence Microscopy

A549 cells were seeded into 6-well plates with 3 mL of media. After 72 hrs, cells (∼50% confluent) were treated with inhibitors. Following incubation, media was aspirated, and cells were treated with inhibitors and 75 nM LysoTracker Red DND-99 (Thermo Fisher # L7528) for 30 min. Media was aspirated and 3 mL of fresh media was added. Invitrogen EVOS M5000 microscope was used to obtain images.

## RESULTS

### Prolonged treatment with CHX shortens 28S and 18S rRNA half-lives

Ribosomes are energetically costly to produce and are highly stable. In fact, ribosomal RNA (rRNA) has been characterized as having half-lives on the order of several days depending on the cell type (26–30). To determine the stability of 28S and 18S rRNA in A549 cells, we used a specific RNA Pol I inhibitor, CX-5461, and assessed half-lives of both 28S and 18S rRNA. Initially, we used 4-thiouridine (4SU) labeling to evaluate the efficiency of CX-5461 to inhibit rRNA synthesis. We observed robust 4SU labeling of 28S and 18S rRNA when cells were treated with vehicle but noticed labeling was completely abolished during treatment with CX-5461, suggesting transcription of new rRNA was sufficiently inhibited (**Figure 1A**). To determine the half-lives of both 28S and 18S rRNA upon treatment with CX-5461, we extracted total RNA from treated cells at six-hour timepoints for 24 hrs and plotted one-phase decay curves. As expected, A549 cells treated with vehicle continued to synthesize new rRNA (**Figure 1B**). In agreement with the literature (26–30), we found that both 28S rRNA and 18S rRNA were stable after treatment with CX-5461 (**Figure 1B**). These half-lives were drastically shortened after prolonged CHX treatment with 28S rRNA having a half-life of approximately 17.91 ± 0.96 hrs and 18S rRNA having a half-life of approximately 23.89 ± 2.44 hrs (**Figure 1B**). This data suggests that ribosomes are stable species and are not regularly targeted for turnover in A549 cells; however, following prolonged treatment with CHX, they become unstable.

**Figure 1:**
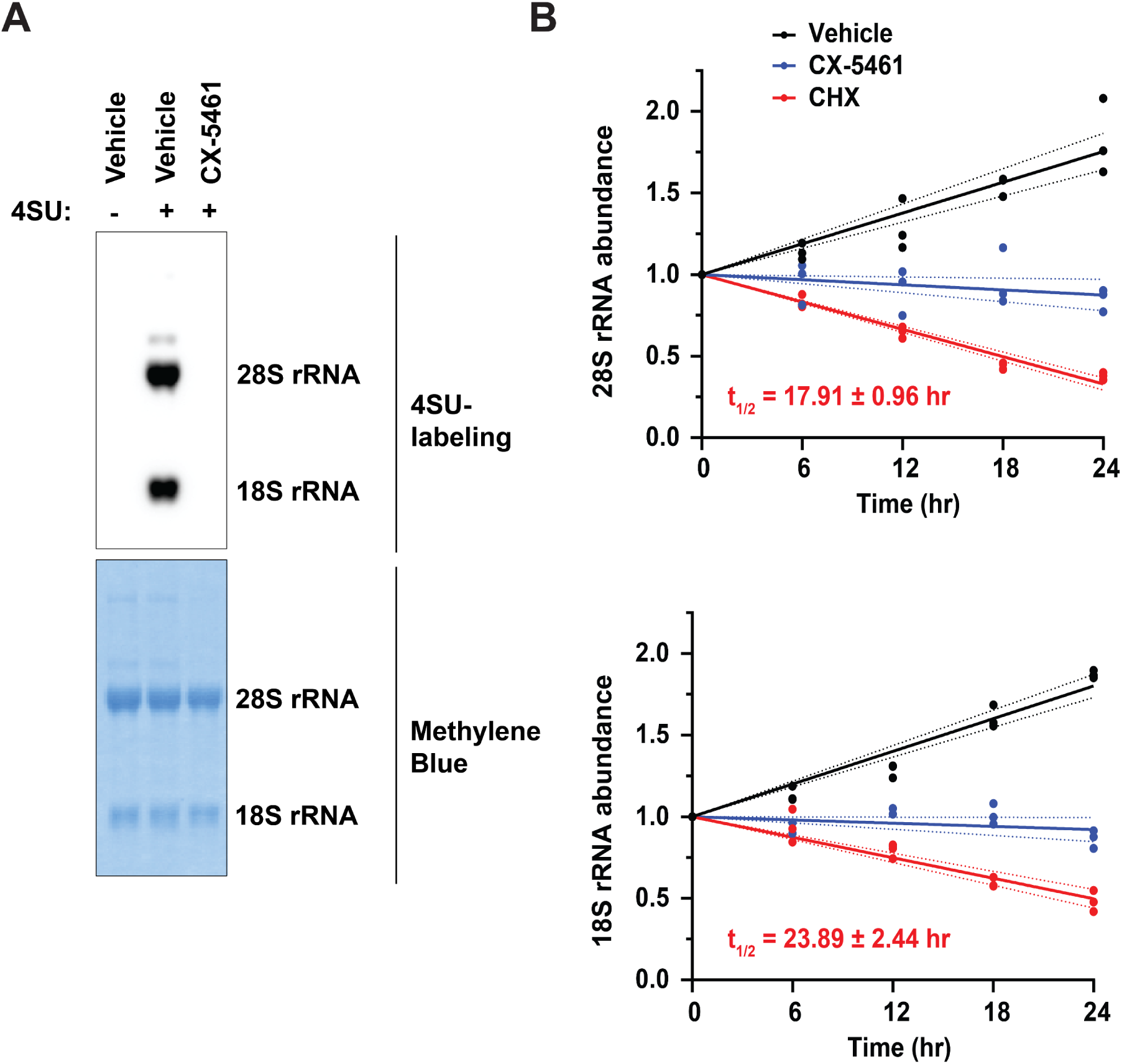
Prolonged treatment with CHX destabilizes 28S and 18S rRNA. A) 4SU-labeling of RNA in the absence or presence of 2.5 μM RNA Pol I inhibitor, CX-5461. Total RNA was stained with methylene blue and served as a loading control. B) 28S and 18S rRNA decay curves from treatment with vehicle (DMSO), 2.5 μM CX-5461 or 100 μg/mL CHX. Total RNA was isolated and analyzed by RNA formaldehyde denaturing gel. Equal volume of total RNA (based on 250 ng RNA for 0 hr control) was loaded. *In vitro* transcribed nLuc (100 ng) was spiked in as a loading control. n=3 biological replicates. A simple linear regression was used to calculate the rRNA half-lives and is shown as the line with the 95% confidence interval included as a watermark. The half-life is reported with the error for the 95% confidence interval range.

### Proteasome inhibitors but not autophagy inhibitors rescue rRNA and ribosomal protein levels upon prolonged translation elongation inhibition

To date, studies have shown that ribosomes are primarily degraded through two decay pathways: autophagy or the ubiquitin-proteasome system (UPS) (31). To test whether ribosomes exposed to prolonged treatments of CHX were being degraded by autophagy or by the UPS, we treated A549 cells with different autophagy inhibitors (SAR405 and BafA1) or different proteasome inhibitors (MG132 and Bortezomib) in the presence of CHX and examined the effects to rRNA levels. Consistent with previous data in HeLa cells (22) and our data (**Figure 1B**), we found that treatment with CHX alone resulted in ∼2-fold reduction of both 28S rRNA and 18S rRNA (**Figure 2A, B**). rRNA levels remained diminished when cells were co-treated with the autophagy inhibitors (**Figure 2A, B**). In contrast, 28S rRNA levels were fully rescued and 18S rRNA levels were partially rescued when cells were co-treated with the proteasome inhibitors (**Figure 2A, B**). We also assessed the impact to 5.8S rRNA and 5S rRNA by denaturing PAGE gel and observed similar depletion and rescue (**Figure 3**). Next, we investigated whether ribosomal proteins were also being degraded by the proteasome. Indeed, when the proteasome was inhibited with MG132 in the presence of CHX, ribosomal proteins associated with the 60S ribosomal subunit (RPL7 and RPL23A) and the 40S ribosomal subunit (RPS3 and RPS6) were either fully or partially rescued (**Figure 2C, D**).

**Figure 2:**
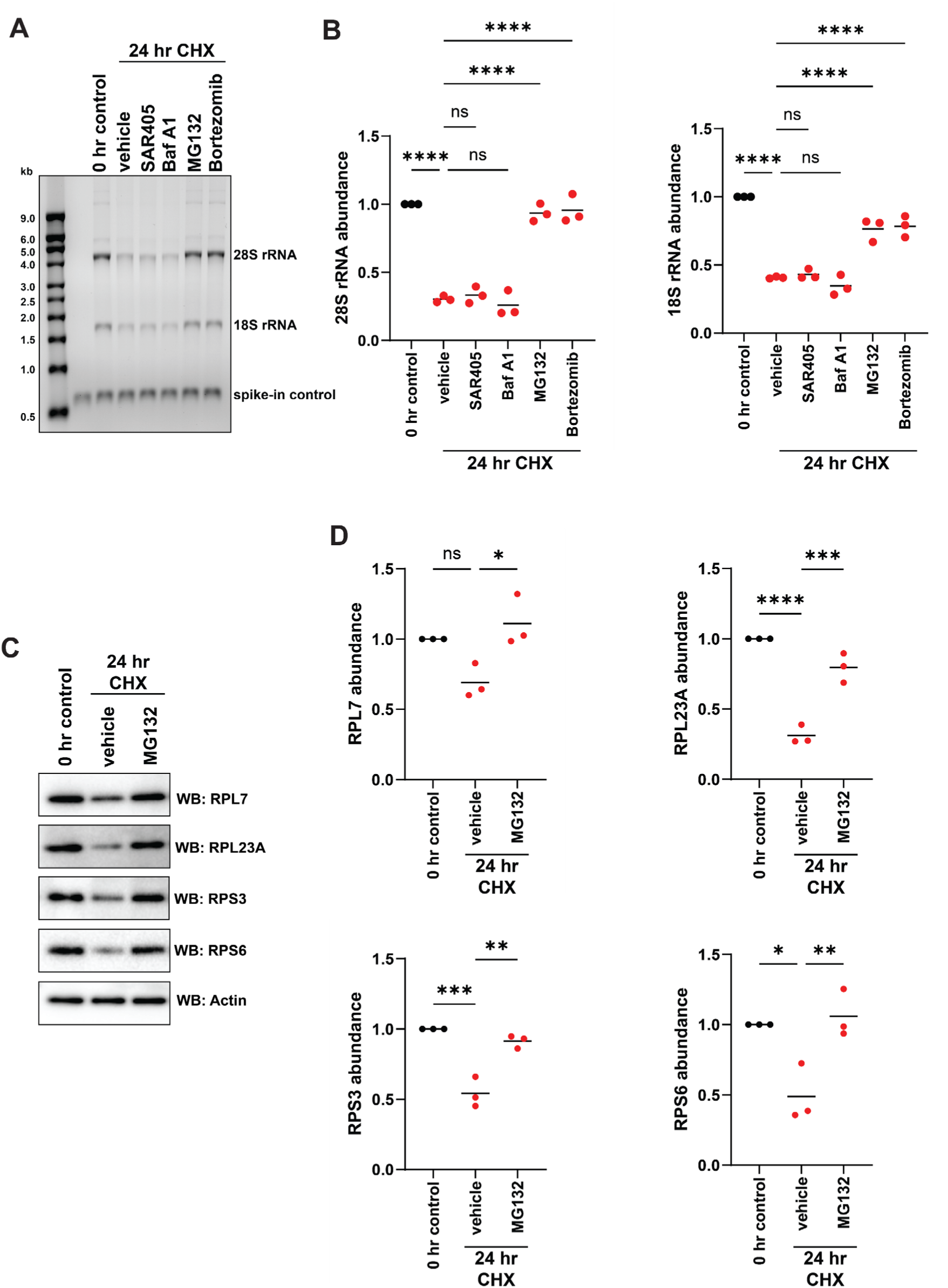
Proteasome inhibitors rescue rRNA and ribosomal protein levels upon prolonged translation inhibition. A) A549 cells were treated for 24 hrs with 100 μg/mL CHX, 100 μg/mL CHX + 10 μM SAR405, 100 μg/mL CHX + 300 nM BafA1, 100 μg/mL CHX + 10 μM MG132, or 100 μg/mL CHX + 10 μM Bortezomib. Total RNA was isolated and analyzed by RNA formaldehyde denaturing gel. Equal volume of total RNA (based on 250 ng RNA for 0 hr control) was loaded. *In vitro* transcribed nLuc (100 ng) was spiked in as a loading control. B) Quantification of 28S and 18S rRNA levels relative to nLuc. Data were set relative to the 0 hr control. Bars represent the mean. n=3 biological replicates. C) Western blot analysis of RPL7, RPL23A, RPS3, and RPS6 protein levels in A549 cells after 24 hr treatment with 100 μg/mL CHX or 100 μg/mL CHX + 10 μM MG132. Actin was used as a loading control. D) Quantification of RPL7, RPL23A, RPS3, and RPS6 protein levels relative to actin protein levels. Data were set relative to the 0 hr control. Bars represent the mean. n=3 biological replicates. Comparisons were made using a one-way ANOVA with Tukey’s multiple comparisons. * = p<0.05. ** = p<0.01. *** = p<0.001. **** = p<0.0001. Exact p-values are reported in **Table 1 and Table 2**.

**Figure 3:**
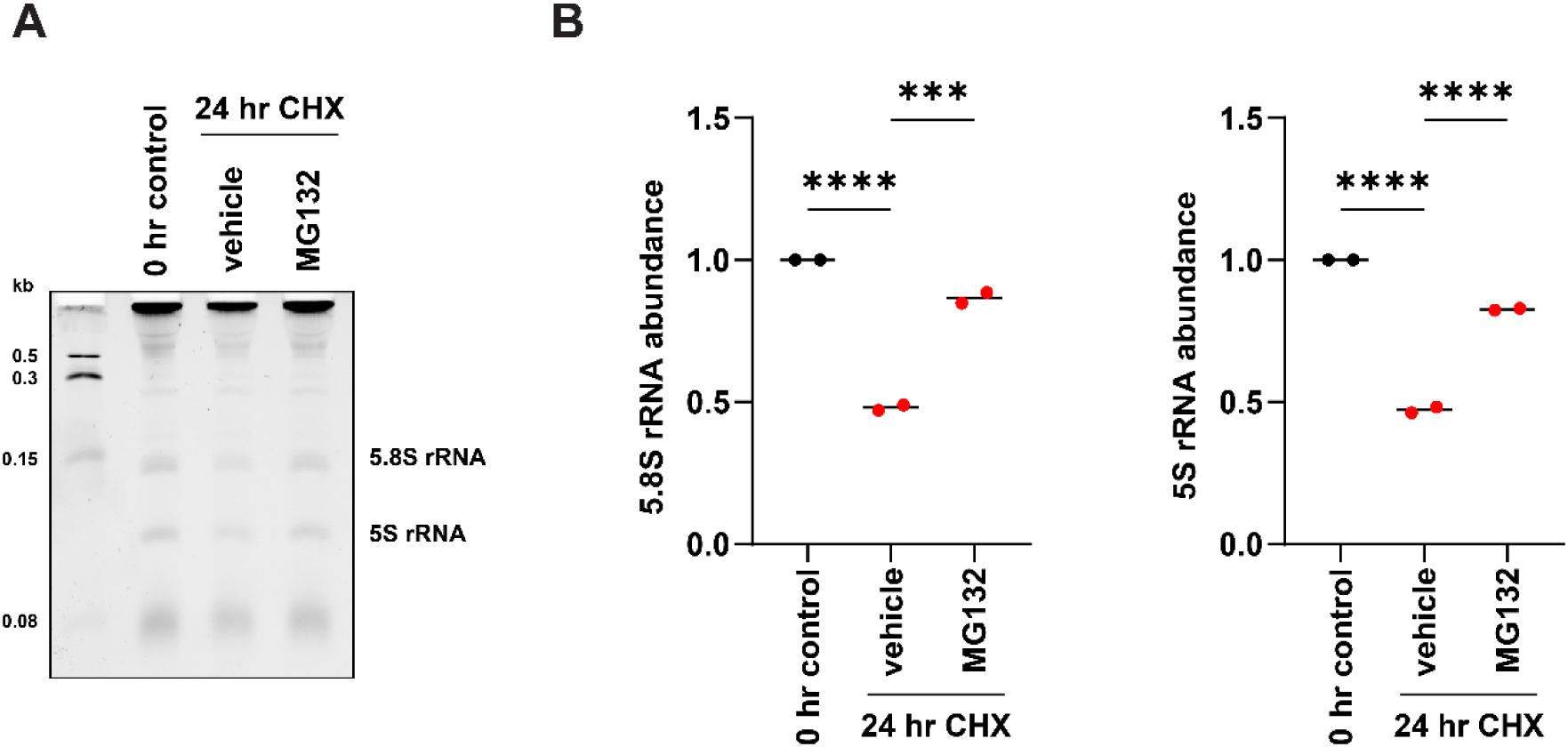
Proteasome inhibitors rescue 5.8S and 5S rRNA levels upon prolonged translation inhibition. A) A549 cells were treated for 24 hrs with 100 μg/mL CHX or 100 μg/mL CHX + 10 μM MG132. Total RNA was isolated and analyzed by PAGE urea denaturing gel. Equal volume of total RNA (based on 1 μg RNA for 0 hr control) was loaded. B) Quantification of 5.8S and 5S rRNA levels. Data were set relative to the 0 hr control. Bars represent the mean. n=2 biological replicates. Comparisons were made using a one-way ANOVA with Tukey’s multiple comparisons. *** = p<0.001. **** = p<0.0001. Exact p-values are reported in **Table 3**.

**Table 1:** P-values of one-way ANOVA with Tukey’s multiple comparisons for. **Figure 2B**

| **28S rRNA abundance** | **p-value** |  |
| --- | --- | --- |
| 0 hr control vs 24 hr CHX | p<0.0001 | **** |
| 24 hr CHX vs 24 hr CHX + SAR405 | p=0.9931 | n.s. |
| 24 hr CHX vs 24 hr CHX + Baf A1 | p=0.9666 | n.s. |
| 24 hr CHX vs 24 hr CHX + MG132 | p<0.0001 | **** |
| 24 hr CHX vs 24 hr CHX + Bortezomib | p<0.0001 | **** |
| **18S rRNA abundance** | **p-value** |  |
| 0 hr control vs 24 hr CHX | p<0.0001 | **** |
| 24 hr CHX vs 24 hr CHX + SAR405 | p=0.9968 | n.s. |
| 24 hr CHX vs 24 hr CHX + Baf A1 | p=0.7657 | n.s. |
| 24 hr CHX vs 24 hr CHX + MG132 | p<0.0001 | **** |
| 24 hr CHX vs 24 hr CHX + Bortezomib | p<0.0001 | **** |

**Table 2:** P-values of one-way ANOVA with Tukey’s multiple comparisons for. **Figure 2D**

| **RPL7 abundance** | **p-value** |  |
| --- | --- | --- |
| 0 hr control vs 24 hr CHX | p=0.0554 | n.s. |
| 24 hr CHX vs 24 hr CHX + MG132 | p=0.0158 | * |
| **RPL23A abundance** | **p-value** |  |
| 0 hr control vs 24 hr CHX | p<0.0001 | **** |
| 24 hr CHX vs 24 hr CHX + MG132 | p=0.0004 | *** |
| **RPS3 abundance** | **p-value** |  |
| 0 hr control vs 24 hr CHX | p=0.0004 | *** |
| 24 hr CHX vs 24 hr CHX + MG132 | p=0.0012 | ** |
| **RPS6 abundance** | **p-value** |  |
| 0 hr control vs 24 hr CHX | p=0.0155 | * |
| 24 hr CHX vs 24 hr CHX + MG132 | p=0.0093 | ** |

**Table 3:** P-values of one-way ANOVA with Tukey’s multiple comparisons for. **Figure 3B**

| **5.8S rRNA abundance** | **p-value** |  |
| --- | --- | --- |
| 0 hr control vs 24 hr CHX | p<0.0001 | **** |
| 24 hr CHX vs 24 hr CHX + MG132 | p=0.0004 | *** |
| **5S rRNA abundance** | **p-value** |  |
| 0 hr control vs 24 hr CHX | p<0.0001 | **** |
| 24 hr CHX vs 24 hr CHX + MG132 | p<0.0001 | **** |

To assess which ribosomal species were being turned over by the proteasome, we used sucrose gradient ultracentrifugation. Again, we observed in the presence of prolonged translation inhibition, a stark reduction in all ribosomal species (40S ribosomal subunits, 60S ribosomal subunits, 80S monosomes, and polysomes) (**Figure 4**) as previously reported in HeLa cells (22). When cells were treated with MG132 in the presence of CHX, a robust accumulation of 80S monosomes and polysomes occurred (**Figure 4**), suggesting that under conditions of prolonged translation elongation inhibition that mature 80S ribosomes are being targeted for proteasome-dependent degradation.

**Figure 4:**
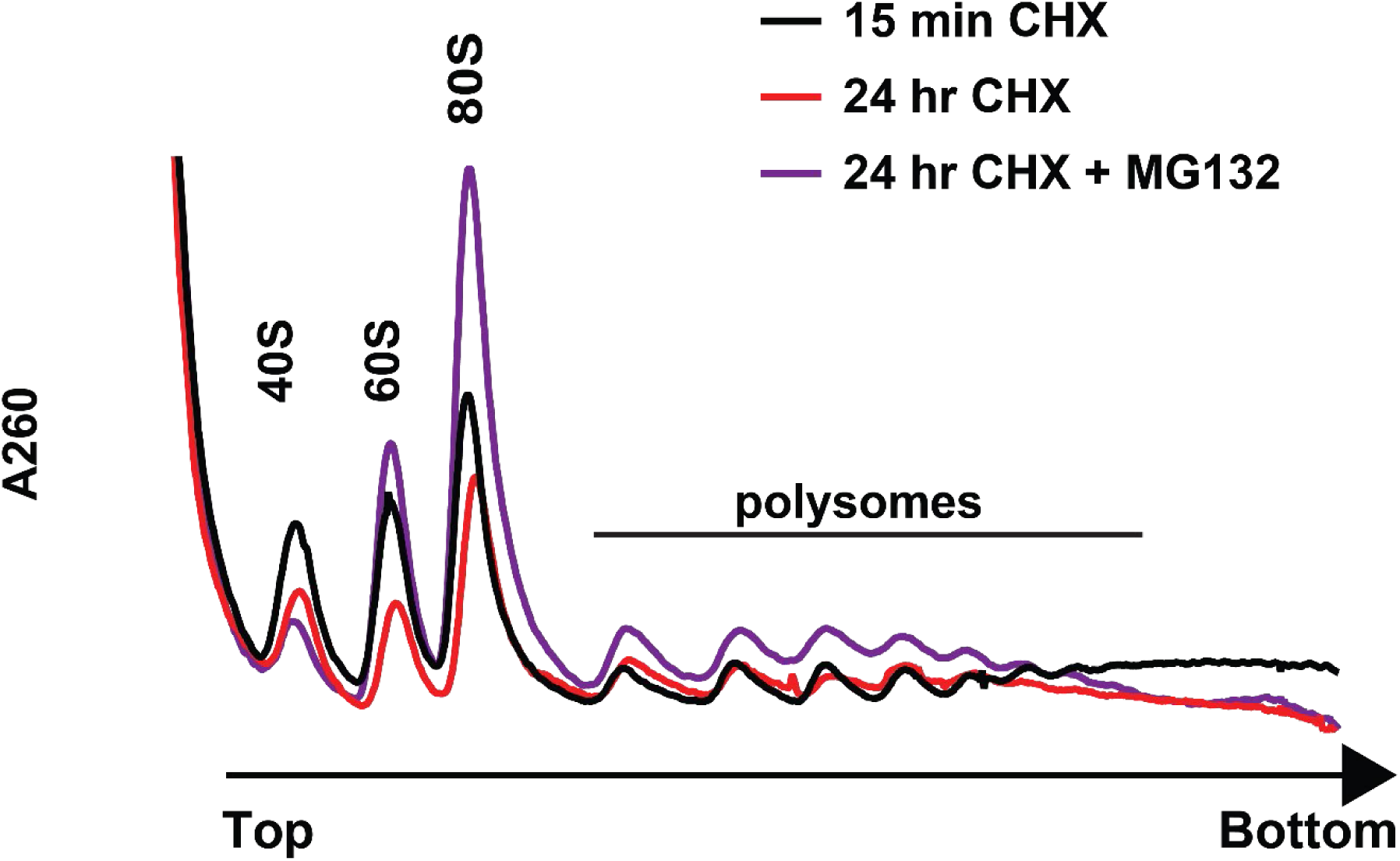
Prolonged co-treatment of CHX with MG132 results in accumulation of 80S monosomes and polysomes in A549 cells. Polysome analysis (10-50% (w/v) sucrose gradients) of A549 cells that had been treated for 15 min with 100 μg/mL CHX (black), 24 hrs with 100 μg/mL CHX (red), or 100 μg/mL CHX + 10 μM MG132 (purple).

### Ubiquitin E1 Inhibitor (TAK243) does not rescue rRNA levels upon prolonged translation elongation inhibition

Typically, proteasome-dependent degradation requires ubiquitin modification to lysine residues on the target protein as a signal to the proteasome that the substrate is available for turnover. Site-specific ubiquitylation of ribosomal proteins has been well characterized for certain ribosome turnover pathways (32–37). Therefore, we next asked whether ubiquitin modification was required for proteasome-dependent turnover of ribosomes in response to prolonged periods of translation elongation inhibition. Ubiquitin is added to the target protein through a relay system that requires an E1 ubiquitin-activating enzyme, an E2 ubiquitin-conjugating enzyme, and an E3 ubiquitin ligase (**Figure 5A**). Consequently, inhibition of the E1 ubiquitin-activating enzyme would prevent downstream ubiquitylation of target proteins. To test this idea, we used an E1 ubiquitin inhibitor, TAK243. To confirm that TAK243 was effective inhibiting the E1 ubiquitin-activating enzyme, recovery of known target proteins and disappearance of known ubiquitin marks was investigated. Initially, cells were pretreated with TAK243 for 1 hr prior to performing co-treatment of TAK243 with CHX for 24 hrs. Cyclin D1 is a known substrate of the UPS (38). During prolonged treatment with CHX alone when no new synthesis of cyclin D1 is occurring, it was observed that the protein is completely degraded in the cell (**Figure 5B**). However, when cells were treated with CHX in the presence of TAK243, cyclin D1 protein levels were fully recovered (**Figure 5B**). Another example of ubiquitylation by E3 ubiquitin ligases is during pre-initiation complex (PIC) collisions. Homoharringtonine (HHT) is a translation inhibitor that binds the A site of the 60S ribosomal subunit and prevents the first aminoacylated-tRNA from entering the ribosome (39,40). This effectively leads to a stalled 80S ribosome on the start codon (41). Trailing PICs scanning for the start codon then collide with the stalled monosome, which elicits a quality control pathway that involves an E3 ubiquitin ligase, RNF10, ubiquitylating RPS2 and RPS3 (33,35,42). We show that after 24 hr treatment with HHT that RPS3 is ubiquitylated as expected (**Figure 5C**). However, during co-treatment with HHT and TAK243, ubiquitylated RPS3 is no longer present (**Figure 5C**).

**Figure 5:**
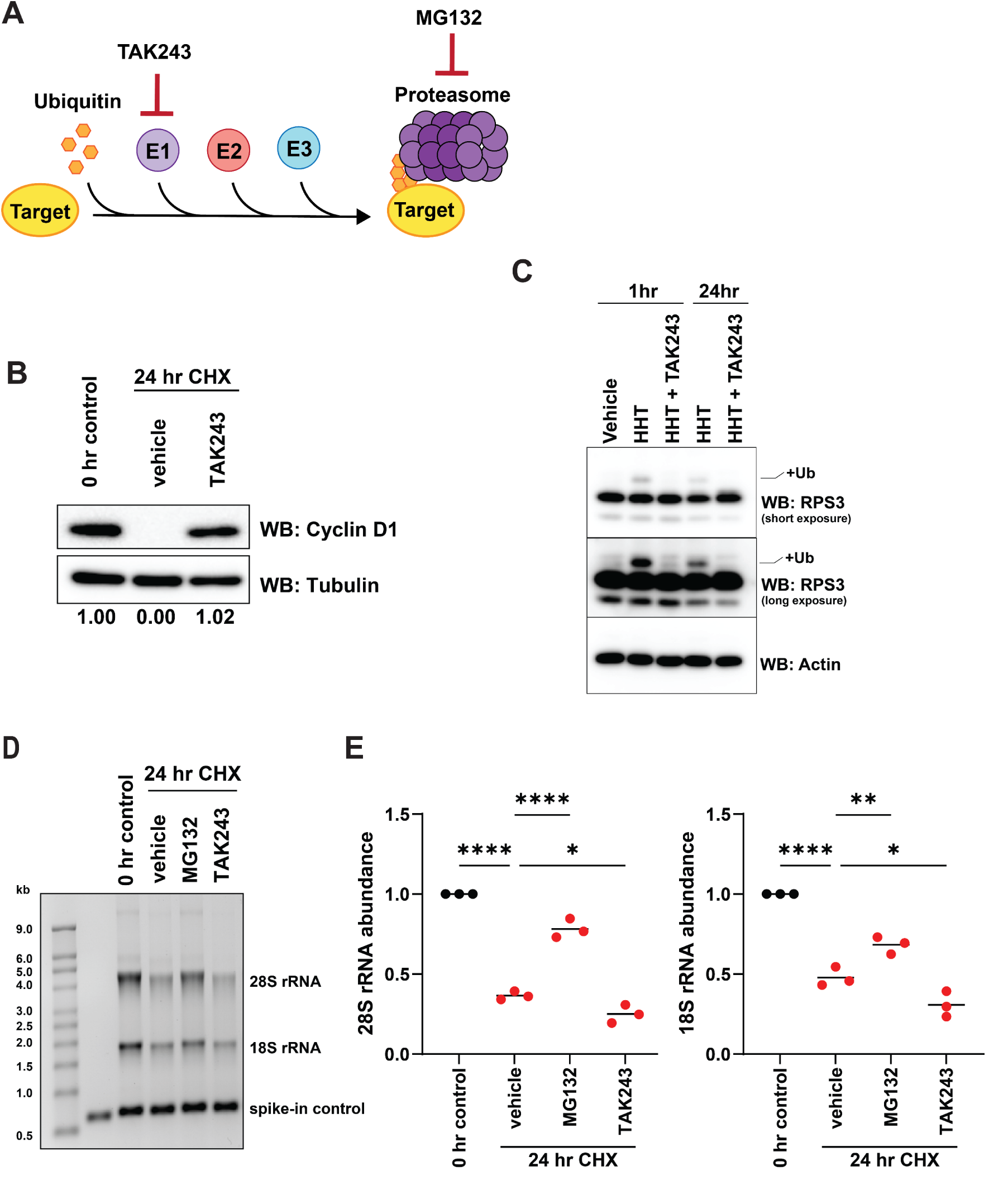
TAK243 does not rescue rRNA upon prolonged translation inhibition. A) Schematic of the ubiquitin-proteasome system (UPS). TAK243 inhibits the E1 ubiquitin-activating enzyme and MG132 inhibits the proteasome. B) Western blot analysis of cyclin D1 protein levels in A549 cells after 24 hr treatment with CHX or CHX + TAK243. A549 cells were pretreated for 1 hr with vehicle (DMSO) or 10 μM TAK243 and then treated for 24 hrs with 100 μg/mL CHX + 50 μM Z-VAD-FMK or 100 μg/mL CHX + 10 μM TAK243 + 50 μM Z-VAD-FMK. Tubulin was used as a loading control. C) Western blot analysis of ubiquitylation of RPS3 in A549 cells after 1 hr or 24 hr treatment with HHT or HHT + TAK243. A549 cells were pretreated for 1 hr with vehicle (DMSO) or 10 μM TAK243 and then treated for 1 hr or 24 hrs with 10 μg/mL HHT + 50 μM Z-VAD-FMK or 10 μg/mL HHT + 10 μM TAK243 + 50 μM Z-VAD-FMK. Actin was used as a loading control. D) A549 cells were pretreated for 1 hr with vehicle (DMSO) or 10 μM TAK243 and then treated for 24 hrs with 100 μg/mL CHX + 50 μM Z-VAD-FMK, 100 μg/mL CHX + 10 μM MG132 + 50 μM Z- VAD-FMK, or 100 μg/mL CHX + 10 μM TAK243 + 50 μM Z-VAD-FMK. Total RNA was isolated and analyzed by RNA formaldehyde denaturing gel. Equal volume of total RNA (based on 250 ng RNA for 0 hr control) was loaded. *In vitro* transcribed nLuc (100 ng) was spiked in as a loading control. E) Quantification of 28S and 18S rRNA levels relative to nLuc. Data were set relative to the 0 hr control. Bars represent the mean. n=3 biological replicates. Comparisons were made using a one-way ANOVA with Tukey’s multiple comparisons. * = p<0.05. ** = p<0.01. **** = p<0.0001. Exact p-values are reported in **Table 4**.

**Table 4:**
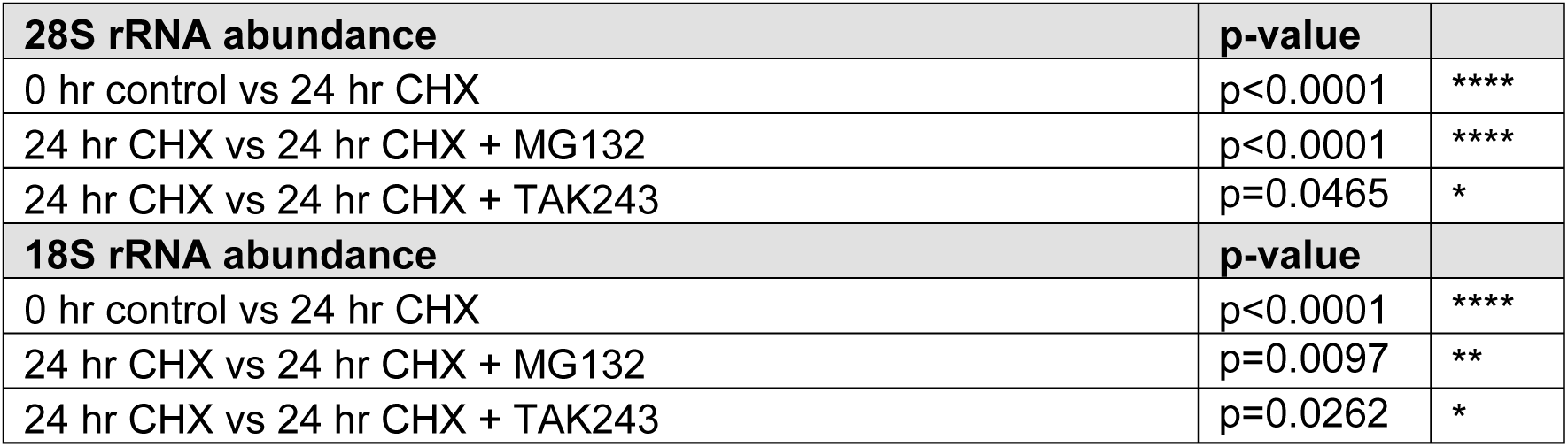
P-values of one-way ANOVA with Tukey’s multiple comparisons for. **Figure 5E**

Knowing that TAK243 could sufficiently inhibit the E1 ubiquitin-activating enzyme in A549 cells, we then focused on its effects to rRNA levels during prolonged CHX treatment. Surprisingly, in contrast to our observation where co-treatment with MG132 rescued rRNA levels, TAK243 co-treatment with CHX showed no recovery (**Figure 5D, E**). This suggests that proteasomal degradation of the ribosome is independent from ubiquitin modification. Additionally, we observed that after prolonged treatment with CHX, that total ubiquitin levels in the cell are depleted (**Figure 6**). Together, this data suggests that a ubiquitin independent mechanism is required to clear ribosomes after extended translation arrest.

**Figure 6:**
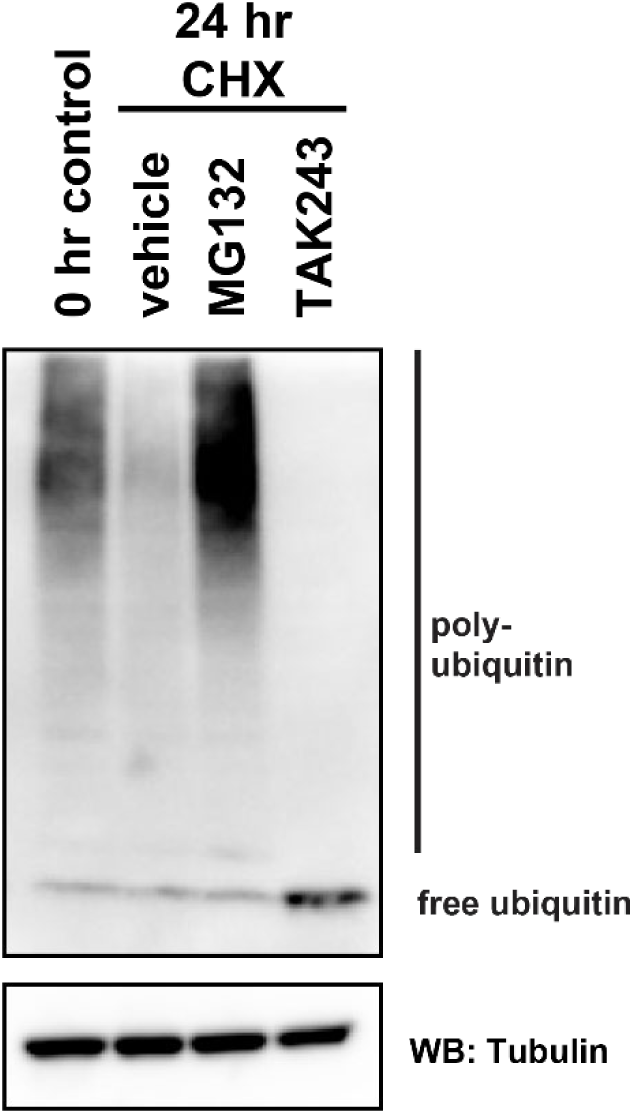
Total ubiquitin protein levels are depleted after 24 hr CHX treatment. Western blot analysis of ubiquitin protein levels in A549 cells after 24 hr treatment with CHX, CHX + MG132, or CHX + TAK243. A549 cells were pretreated for 1 hr with vehicle (DMSO) or 10 μM TAK243 and then treated for 24 hrs with 100 μg/mL CHX + 50 μM Z-VAD-FMK, 100 μg/mL CHX + 10 μM MG132 + 50 μM Z-VAD-FMK, or 100 μg/mL CHX + 10 μM TAK243 + 50 μM Z-VAD-FMK. Tubulin was used as a loading control.

### Ribosome degradation is stimulated by stalled monosomes in different states on the mRNA

To assess if ribosomes trapped in different states on the mRNA were also targeted for degradation, we treated A549 cells with prolonged high doses of several other translation elongation inhibitors: anisomycin (ANS), homoharringtonine (HHT), and lactimidomycin (LTM). In brief, ANS binds to the A site of 60S ribosomal subunit and prevents peptidyl transferase activity (43) while LTM binds to the E site of the 60S ribosomal subunit and blocks the first translocation cycle (21). Treatment with ANS and CHX result in ribosomes trapped along the ORF with occupied A sites whereas treatment with HHT and LTM generate stalled ribosomes at the start codon with empty or occupied A-sites, respectively. We observed a similar ∼2-fold reduction of both 28S rRNA and 18S rRNA levels for all translation elongation inhibitors as was previously observed with CHX treatment (**Figure 7**). Co-treatment with MG132 was able to fully or partially rescue 28S rRNA and 18S rRNA levels (**Figure 7**). This data suggests that this ribosome turnover pathway can recognize ribosomes trapped in different states and conformations on the mRNA.

**Figure 7:**
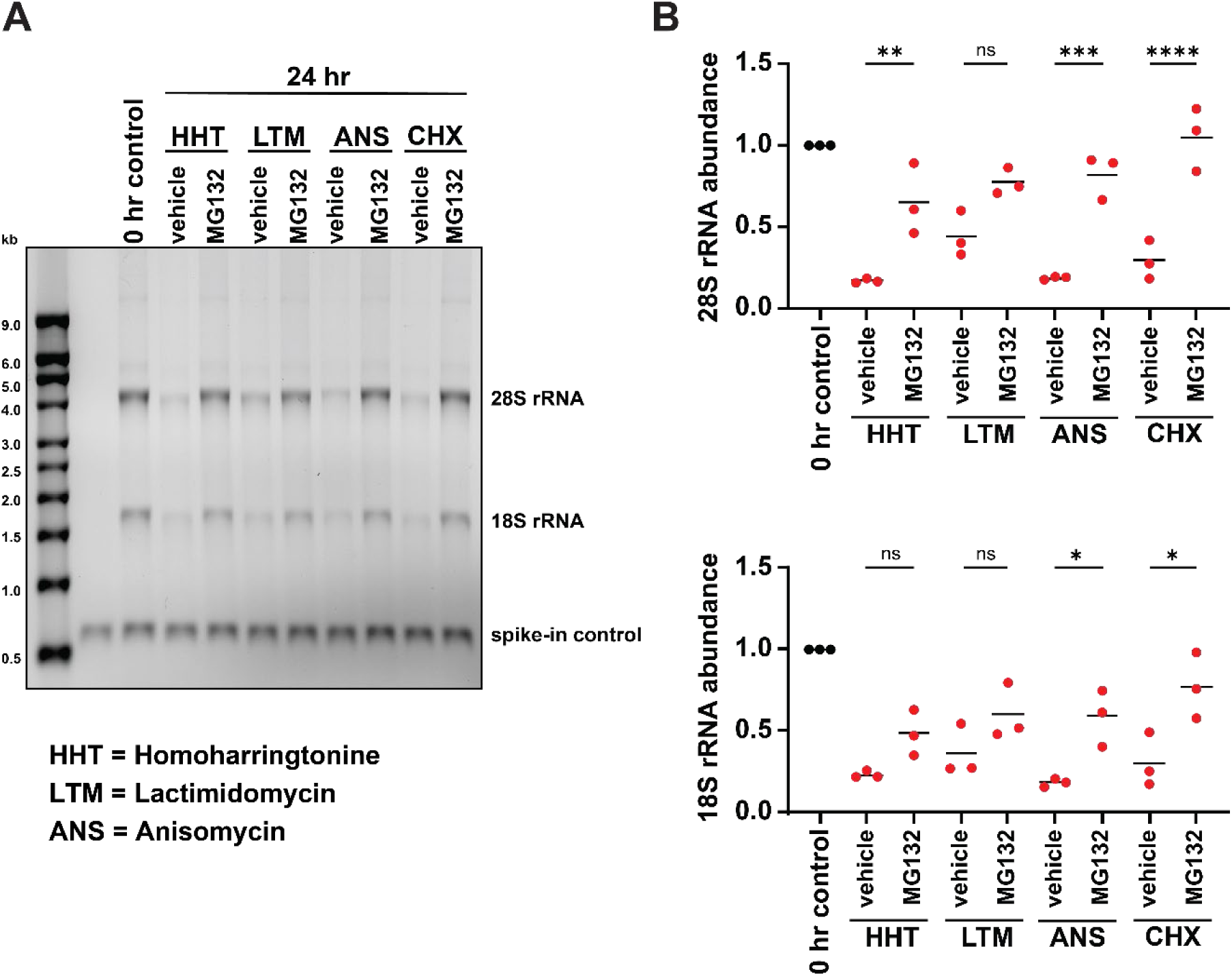
Ribosomes trapped on the mRNA in different states are indiscriminately targeted for proteasomal degradation. A) A549 cells were treated for 24 hrs with 10 μg/mL HHT, 10 μg/mL HHT + 10 μM MG132, 5 μM LTM, 5 μM LTM + 10 μM MG132, 10 μg/mL ANS, 10 μg/mL ANS + 10 μM MG132, 100 μg/mL CHX, or 100 μg/mL CHX + 10 μM MG132. Total RNA was isolated and analyzed by RNA formaldehyde denaturing gel. Equal volume of total RNA (based on 250 ng RNA for 0 hr control) was loaded. *In vitro* transcribed nLuc (100 ng) was spiked in as a loading control. B) Quantification of 28S and 18S rRNA levels relative to nLuc. Data were set relative to the 0 hr control. Bars represent the mean. n=3 biological replicates. Comparisons were made using a one-way ANOVA with Tukey’s multiple comparisons. * = p<0.05. ** = p<0.01. *** = p<0.001. **** = p<0.0001. Exact p-values are reported in **Table 5**.

**Table 5:** P-values of one-way ANOVA with Tukey’s multiple comparisons for. **Figure 7B**

| **28S rRNA abundance** | **p-value** |  |
| --- | --- | --- |
| 24 hr HHT vs 24 hr HHT + MG132 | p=0.0045 | ** |
| 24 hr LTM vs 24 hr LTM + MG132 | p=0.0932 | n.s. |
| 24 hr ANS vs 24 hr ANS + MG132 | p=0.0002 | *** |
| 24 hr CHX vs 24 hr CHX + MG132 | p<0.0001 | **** |
| **18S rRNA abundance** | **p-value** |  |
| 24 hr HHT vs 24 hr HHT + MG132 | p=0.4312 | n.s. |
| 24 hr LTM vs 24 hr LTM + MG132 | p=0.5147 | n.s. |
| 24 hr ANS vs 24 hr ANS + MG132 | p=0.0430 | * |
| 24 hr CHX vs 24 hr CHX + MG132 | p=0.0147 | * |

Considering that ribosome collisions are common recognition platforms to many ribosome quality control pathways, we next asked whether prolonged treatment with CHX resulted in collided disomes. To test whether stalled monosomes or collided disomes were present following prolonged treatment with CHX, we treated A549 cells with MG132 to preserve the targeted ribosome complexes and then treated the cell lysate with RNase and investigated the presence of nuclease-resistant disome peaks. As a negative control, we treated cells with CHX and MG132 for 15 min and, as expected, observed the presence of an 80S monosome peak but no disome peak (**Figure 8**). A similar phenotype for cells treated with CHX and MG132 for 24 hrs resulted (**Figure 8**) suggesting that stalled monosomes are present following prolonged treatment with CHX and are therefore the ribosome species being targeted by the proteasomal-dependent degradation pathway.

**Figure 8:**
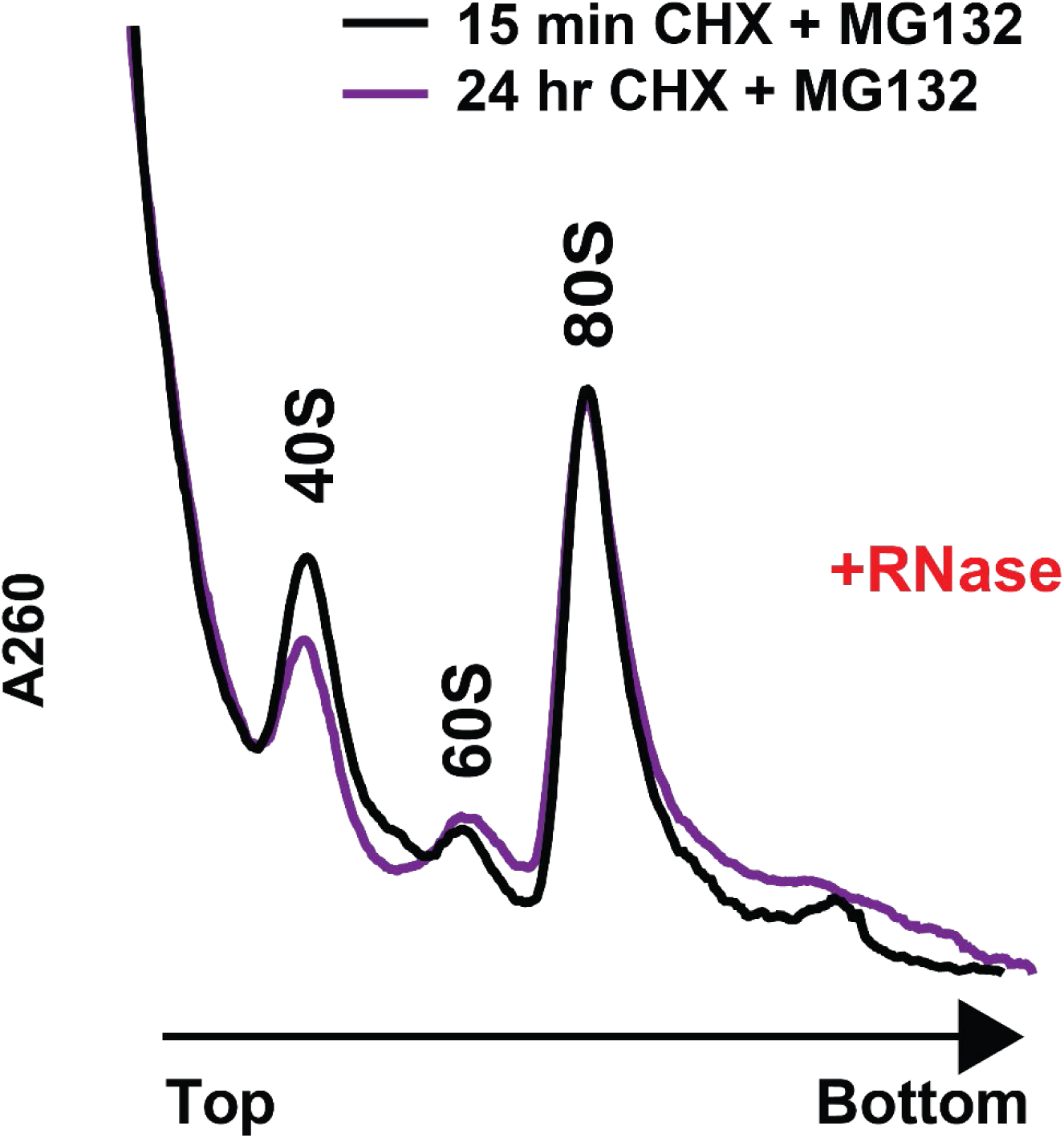
Ribosome turnover following prolonged treatment with elongation inhibitors occurs independently of ribosome collisions. Polysome analysis (10-50% (w/v) sucrose gradients) of the S7 micrococcal nuclease (RNase)-treated lysates from A549 cells that had been treated for 15 min with 100 μg/mL CHX + 10 μM MG132 (black) or 24 hrs with 100 μg/mL CHX + 10 μM MG132 (purple).

### Inhibition of ZAK and the RSR does not protect ribosomes from degradation after prolonged periods of CHX treatment

It has previously been shown that stalled 80S monosomes initiate the ribotoxic stress response (RSR) pathway. During 80S ribosome stalling from short exposure to elongation inhibitors, it was observed that ZAKα becomes phosphorylated and activated (19). This leads to downstream phosphorylation and activation of mitogen-activated protein (MAP) kinases, p38 and JNK, which govern signaling cascades that result in cell cycle arrest and cell apoptosis, respectively (17–20). To investigate whether prolonged treatment with high dosages of CHX results in activation of the RSR pathway and if inactivation of the RSR pathway could rescue rRNA levels, we performed siRNA knockdown of ZAK or inhibited ZAK activity and examined effects to rRNA levels. Initially, we confirmed that ZAK was efficiently depleted in A549 cells following siRNA knockdown (**Figure 9A**). Despite almost complete reduction of ZAK protein levels, we still observed some activation of p38 (as evidenced by phospho-p38 levels) after treatment with CHX for 24 hrs (**Figure 9A**). When 28S rRNA and 18S rRNA were assessed under ZAK knockdown conditions, we observed no rescue of rRNA levels (**Figure 9B, C**). To corroborate these results, we also performed similar experiments with a ZAK inhibitor (ZAKi), PLX4720 (44). A549 cells were pretreated with ZAKi for 1 hr before performing co-treatment of ZAKi with CHX for 24 hrs. Again, despite high dosages of ZAKi, some activation of p38 was still observed (**Figure 9D**) and 28S rRNA and 18S rRNA levels were unable to be recovered (**Figure 9E, F**). Although this data is suggestive that prolonged treatments with high dosages of CHX does not activate a ZAK-dependent RSR pathway, given that there were still some elevated levels of phopsoho-p38 under these conditions, it is still entirely possible that signaling downstream of p38 activation is important for the observed mature ribosome degradation.

**Figure 9:**
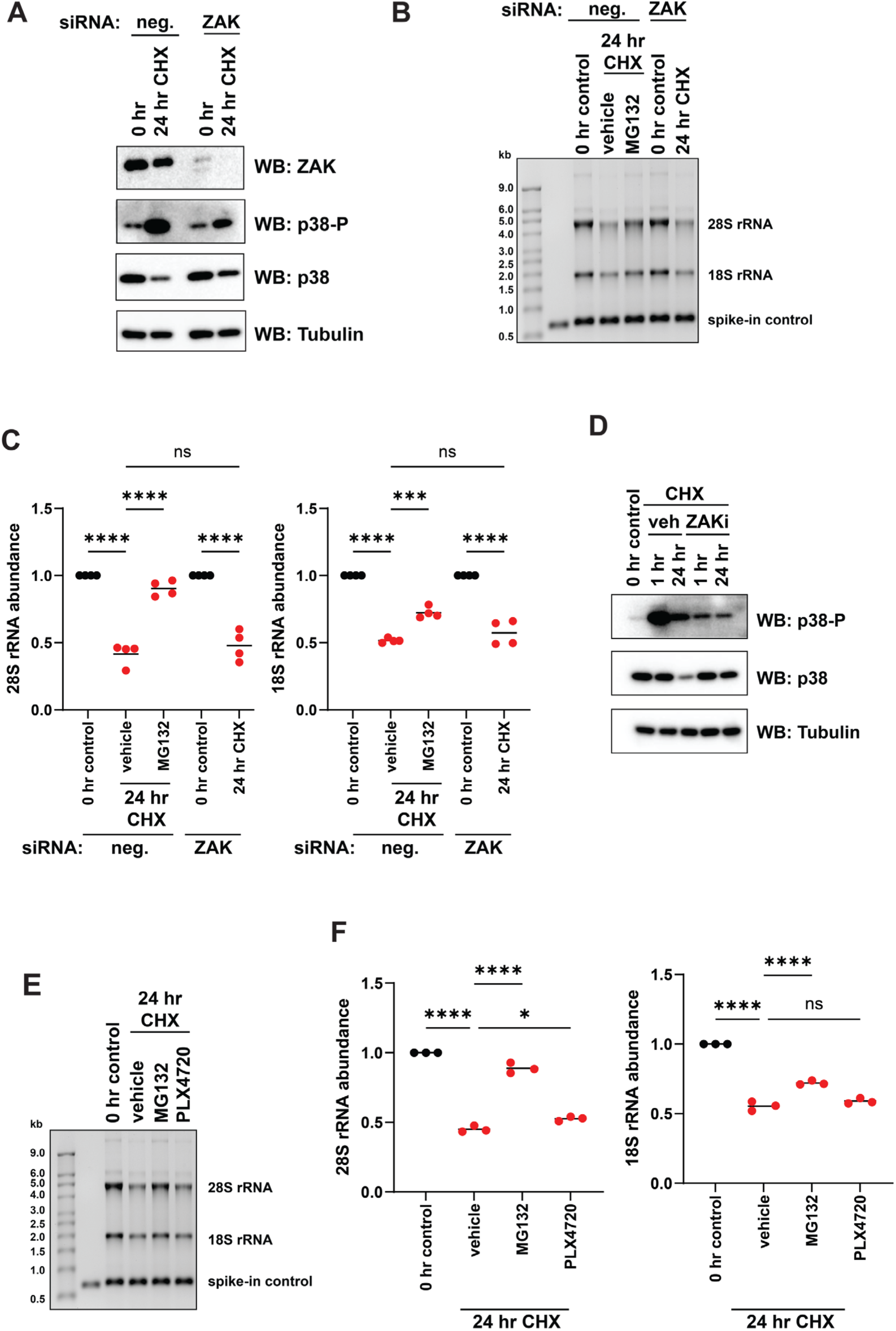
Inhibition of ZAK does not protect ribosomes from degradation after prolonged periods of CHX treatment. A) Western blot analysis of p38 and p38-P protein levels in A549 cells after 24 hr treatment with 100 μg/mL CHX in the presence or absence of ZAK. Negative (neg.) #1 and ZAK #1 targeting siRNAs were used. Tubulin was used as a loading control. B) A549 cells were treated for 24 hrs with 100 μg/mL CHX or 100 μg/mL CHX + 10 μM MG132 in the presence or absence of ZAK. Total RNA was isolated and analyzed by RNA formaldehyde denaturing gel. Equal volume of total RNA (based on 250 ng RNA for 0 hr control) was loaded. *In vitro* transcribed nLuc (100 ng) was spiked in as a loading control. C) Quantification of 28S and 18S rRNA levels relative to nLuc. Data were set relative to the 0 hr control. Bars represent the mean. n=4 biological replicates. D) Western blot analysis of p38 and p38-P protein levels in A549 cells after 1 hr or 24 hr treatment with 100 μg/mL CHX ± 100 μM PLX4720 (ZAKi). A549 cells were pretreated for 1 hr with vehicle (DMSO) or 100 μM ZAKi prior to treatment. Tubulin was used as a loading control. E) A549 cells were pretreated for 1 hr with vehicle (DMSO) or 100 μM ZAKi and then treated with 100 μg/mL CHX, 100 μg/mL CHX + 10 μM MG132, or 100 μg/mL CHX + 100 μM (ZAKi). Total RNA was isolated and analyzed by RNA formaldehyde denaturing gel. Equal volume of total RNA (based on 250 ng RNA for 0 hr control) was loaded. *In vitro* transcribed nLuc (100 ng) was spiked in as a loading control. F) Quantification of 28S and 18S rRNA levels relative to nLuc. Data were set relative to the 0 hr control. Bars represent the mean. n=3 biological replicates. Comparisons were made using a one-way ANOVA with Tukey’s multiple comparisons. * = p<0.05. *** = p<0.001. **** = p<0.0001. Exact p-values are reported in **Table 6 and Table 7**.

**Table 6:** P-values of one-way ANOVA with Tukey’s multiple comparisons for. **Figure 9C**

| **28S rRNA abundance** | **p-value** |  |
| --- | --- | --- |
| 0 hr control (neg.) vs 24 hr CHX (neg.) | p<0.0001 | **** |
| 24 hr CHX (neg.) vs + 24 hr CHX + MG132 (neg.) | p<0.0001 | **** |
| 0 hr control (ZAK) vs 24 hr CHX (ZAK) | p<0.0001 | **** |
| 24 hr CHX (neg.) vs 24 hr CHX (ZAK) | p=0.6888 | n.s. |
| **18S rRNA abundance** | **p-value** |  |
| 0 hr control (neg.) vs 24 hr CHX (neg.) | p<0.0001 | **** |
| 24 hr CHX (neg.) vs + 24 hr CHX + MG132 (neg.) | p=0.0001 | *** |
| 0 hr control (ZAK) vs 24 hr CHX (ZAK) | p<0.0001 | **** |
| 24 hr CHX (neg.) vs 24 hr CHX (ZAK) | p=0.4239 | n.s. |

**Table 7:** P-values of one-way ANOVA with Tukey’s multiple comparisons for. **Figure 9F**

| **28S rRNA abundance** | **p-value** |  |
| --- | --- | --- |
| 0 hr control vs 24 hr CHX | p<0.0001 | **** |
| 24 hr CHX vs 24 hr CHX + MG132 | p<0.0001 | **** |
| 24 hr CHX vs 24 hr CHX + PLX4720 | p=0.0171 | * |
| **18S rRNA abundance** | **p-value** |  |
| 0 hr control vs 24 hr CHX | p<0.0001 | **** |
| 24 hr CHX vs 24 hr CHX + MG132 | p<0.0001 | **** |
| 24 hr CHX vs 24 hr CHX + PLX4720 | p=0.2311 | n.s. |

### Free 80S ribosomes are also subject to ribosome turnover

It has previously been reported that when cells are exposed to heat stress the proteasome can be recruited with the assistance of heat shock proteins to improperly folded nascent polypeptides for targeted degradation (45). During translational arrest of ribosomes along the mRNA with translation elongation inhibitors, the incomplete nascent polypeptide protruding out of the ribosome is unlikely to adopt its correct fold. Therefore, we next asked whether the nascent polypeptide was important for recognition by the degradation pathway. To test for this, we used puromycin, an aminoacyl-tRNA analog, that becomes incorporated into the nascent polypeptide during active elongation and leads to chain termination and dissociation of the 80S ribosome from the mRNA (46,47). Although some studies argue that pretreatment with translation elongation inhibitors prevents puromycylation of the nascent polypeptide, others have shown that puromycylation can still occur under these conditions and that ribosomes remain bound to the mRNA (48,49). As such, we initially pretreated A549 cells with CHX for 20 min and then performed co-treatment of CHX and puromycin for 24 hrs. We once again observed an approximate ∼2-fold reduction of both 28S rRNA and 18S rRNA levels (**Figure 10**). Interestingly, we also observed similar levels of depletion of 28S rRNA and 18S rRNA after prolonged treatment with just puromycin (**Figure 10**). This data suggest that the nascent polypeptide is dispensable for recognition and that free 80S ribosomes can also be targeted for degradation.

**Figure 10:**
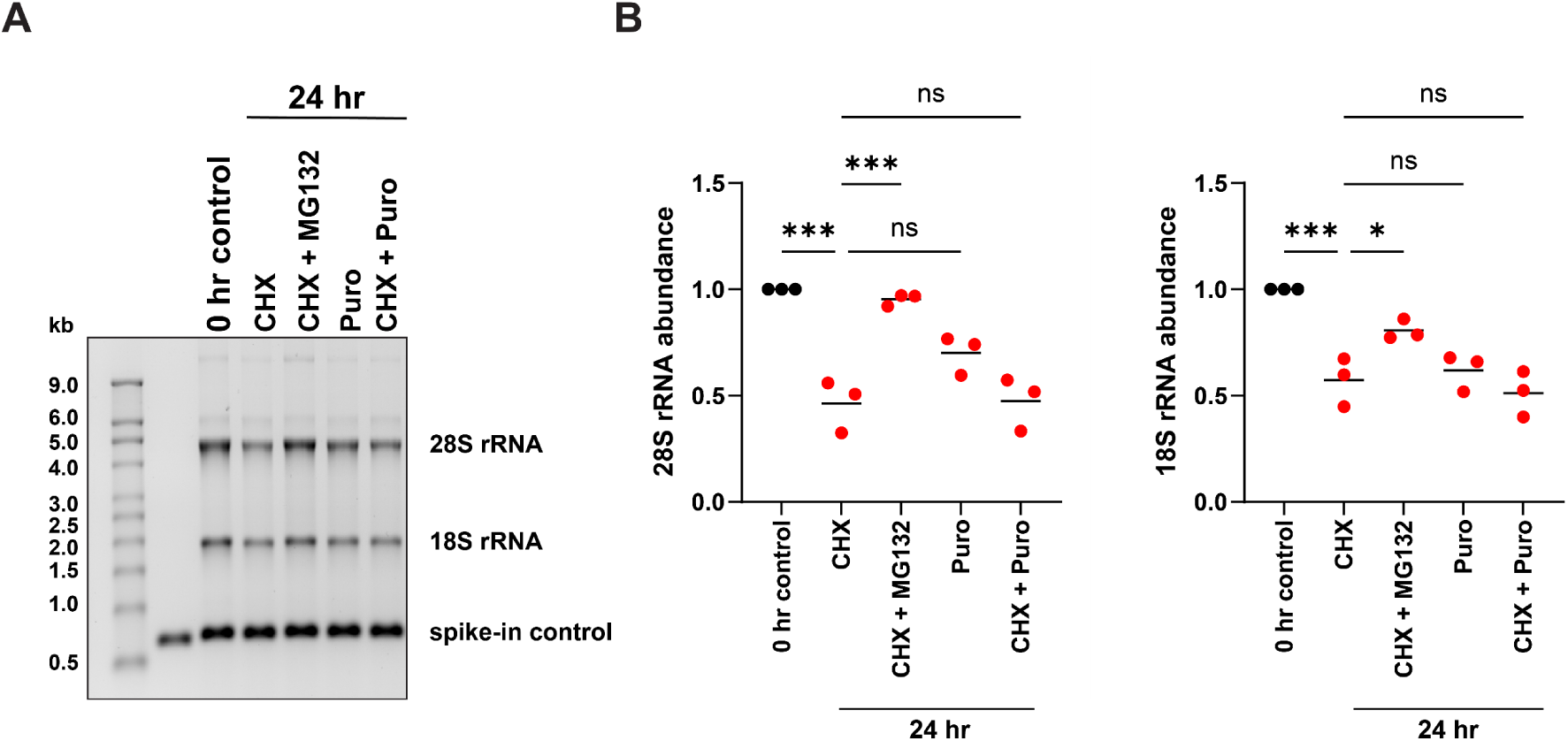
Dissociation of 80S ribosomes from mRNAs also causes ribosome turnover. A) A549 cells were pretreated with vehicle (DMSO) or 100 μg/mL CHX for 20 min and then treated for 24 hrs with 100 μg/mL CHX + 50 μM Z-VAD-FMK, 100 μg/mL CHX + 10 μM MG132 + 50 μM Z-VAD-FMK, 333 μg/mL puromycin (Puro) + 50 μM Z-VAD-FMK, or 100 μg/mL CHX + 333 μg/mL Puro + 50 μM Z-VAD-FMK. Total RNA was isolated and analyzed by RNA formaldehyde denaturing gel. Equal volume of total RNA (based on 250 ng RNA for 0 hr control) was loaded. *In vitro* transcribed nLuc (100 ng) was spiked in as a loading control. B) Quantification of 28S and 18S rRNA levels relative to nLuc. Data were set relative to the 0 hr control. Bars represent the mean. n=3 biological replicates. Comparisons were made using a one-way ANOVA with Tukey’s multiple comparisons. * = p<0.05. *** = p<0.001. Exact p-values are reported in **Table 8**.

**Table 8:** P-values of one-way ANOVA with Tukey’s multiple comparisons for. **Figure 10B**

| **28S rRNA abundance** | **p-value** |  |
| --- | --- | --- |
| 0 hr control vs 24 hr CHX | p=0.0002 | *** |
| 24 hr CHX vs 24 hr CHX + MG132 | p=0.0004 | *** |
| 24 hr CHX vs 24 hr PURO | p=0.0552 | n.s. |
| 24 hr CHX vs 24 hr CHX + PURO | p=0.9999 | n.s. |
| **18S rRNA abundance** | **p-value** |  |
| 0 hr control vs 24 hr CHX | p=0.0007 | *** |
| 24 hr CHX vs 24 hr CHX + MG132 | p=0.0396 | * |
| 24 hr CHX vs 24 hr PURO | p=0.9590 | n.s. |
| 24 hr CHX vs 24 hr CHX + PURO | p=0.8903 | n.s. |

### Lysosome leakage phenocopies rRNA degradation

Lysosomes contain specific RNases and proteases that are normally sequestered from the cytosol and, consequently, under physiological conditions cannot degrade ribosomes. However, under conditions that could lead to lysosomal membrane permeabilization (LMP), the release of these factors into the cytoplasm could result in ribosome turnover. Previous studies have indicated that components in the lysosome are able to degrade ribosomes in a process referred to as RNautophagy (50). Therefore, we next asked whether disruption of lysosome membrane integrity was important for global ribosome degradation. To permeabilize the lysosome membrane, A549 cells were treated with L-leucyl-L-leucine methyl ester (hydrochloride) (LLOMe). Upon being endocytosed by the lysosome, LLOMe is converted into poly-leucine peptides which lead to lysis of the membrane (51,52). It has been shown following membrane permeabilization that lysosomal proteases, such as N-acetyl-β-d-glucosaminidase (NAG) and cathepsin B, are able to leak out into the cytosol (51). We observed after treating A549 cells with LLOMe for 8 hr that 28S rRNA and 18S rRNA levels phenocopied that of 24 hr CHX treatment (**Figure 11A, B**). To further examine if lysosome permeabilization was responsible for ribosome turnover observed after prolonged treatments with translation elongation inhibitors, we measured LMP using LysoTracker Red, a fluorescent dye that stains the acidic compartment of lysosomes. When A549 cells were untreated, a robust accumulation of signal was observed in lysosomes (**Figure 11C**).

**Figure 11:**
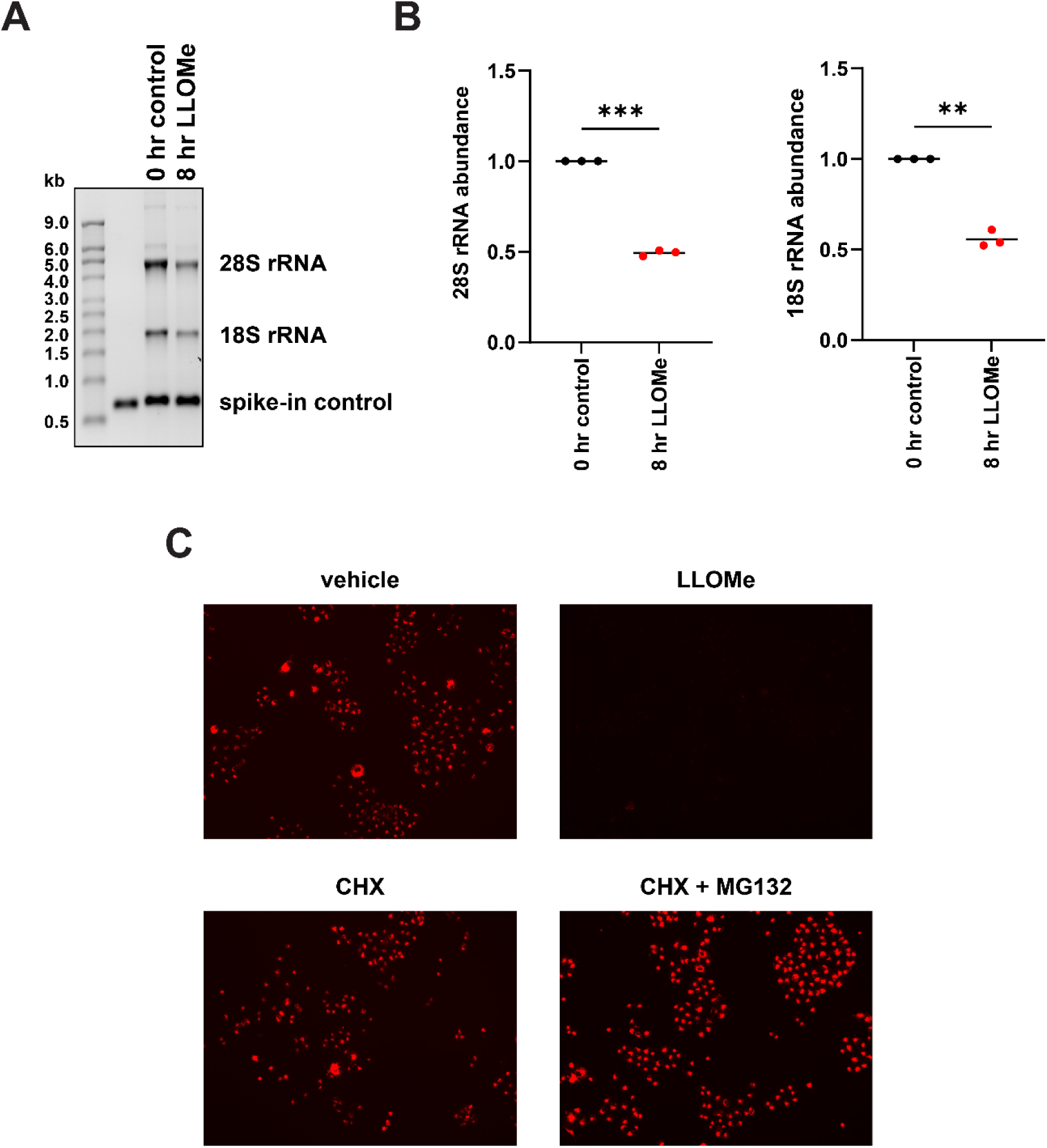
Lysosome permeabilization is not responsible for ribosome degradation after prolonged treatment with translation elongation inhibitors. A) A549 cells were treated for 8 hr with 5 mM L-Leucyl-L-Leucine methyl ester (LLOMe). Total RNA was isolated and analyzed by RNA formaldehyde denaturing gel. Equal volume of total RNA (based on 250 ng RNA for 0 hr control) was loaded. *In vitro* transcribed nLuc (100 ng) was spiked in as a loading control. B) Quantification of 28S and 18S rRNA levels relative to nLuc. Data were set relative to the 0 hr control. Bars represent the mean. n=3 biological replicates. C) Epifluorescence images of A549 cells treated for either 8 hrs with vehicle (DMSO) or 5mM LLOMe or 24 hrs with 100 μg/mL CHX or 100 μg/mL CHX + 10 μM MG132. Lysosomes were stained with 75 nM LysoTracker Red DND-99 for 30 min. Comparisons were made using a two-tailed unpaired t-test with Welch’s correction. ** = p<0.01. *** = p<0.001. Exact p-values are reported in **Table 9**.

**Table 9:**
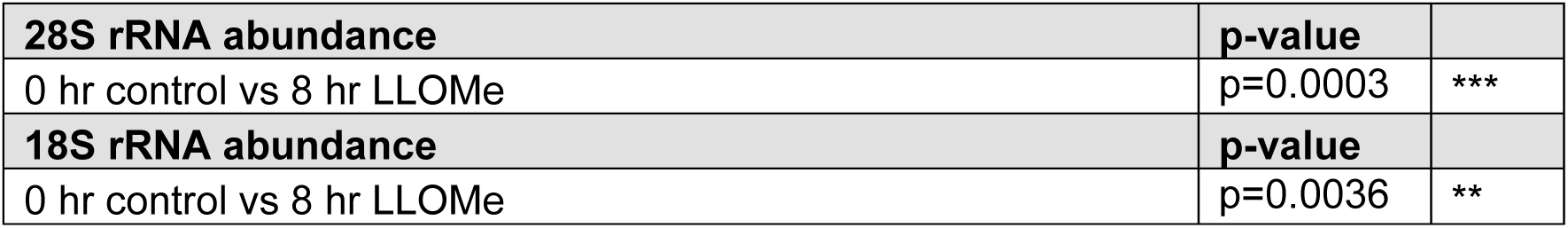
P-values of two-tailed unpaired t tests with Welch’s correction for. **Figure 11B**

However, when the cells were treated with LLOMe for 8 hr, the signal was greatly dissipated (**Figure 11C**). A549 cells treated for 24 hrs with CHX produced a similar signal intensity as vehicle (**Figure 11C**). This suggests that increased LMP is not occurring after treatment with CHX. It was also observed that cells treated with CHX and MG132 for 24 hrs had a higher overall signal intensity compared to vehicle (**Figure 11C**). This agrees with another study in astrocytes that showed that prolonged treatment with MG132 resulted in increased acidic compartments (53).

## DISCUSSION

RQC pathways are critical response mechanisms used by the cell to degrade otherwise detrimental proteins and mRNAs. These pathways take advantage of the ribosomes processivity along the mRNA to survey for abnormalities. Due to the one-directional nature of translation, when ribosome encounter obstructions in the mRNA, they stall and generally lead to ribosome collisions. These collisions act as beacons to the cell that an issue exists, and, consequently, the necessary factors are recruited to the site to resolve the defect in elongation. Importantly, outside the scope of specialized cases, in most instances the ribosomal subunits are recycled off the mRNA so they can be used in future rounds of translation. Given that ribosomes are energetically costly to produce (54), it stands to reason that RQC mechanisms have evolved to primarily target and remove the nascent polypeptide and the defective mRNA while salvaging the ribosomal subunits.

In contrast with the canonical RQC pathway, we show that ribosomes stalled following prolonged translation elongation inhibition with high concentrations of CHX are not recycled but rather degraded (**Figure 2**). Our observations suggest a quality control pathway that further deviates from known RQC pathways as conventional recognition hallmarks such as ribosome collisions are absent (**Figure 8**) and site-specific ubiquitylation of RPs appears nonessential (**Figure 5**). Additionally, we show that ribosomes trapped in different states on the mRNA (**Figure 7**), ribosomes trapped on the mRNA without the nascent polypeptide (**Figure 10**) and even free 80S ribosomes are all targeted for degradation (**Figure 10**). This data argues that a bulk degradation pathway such as autophagy would likely be required to recognize these globally stalled ribosomes. Interestingly, we did not observe rescue of 28S rRNA and 18S rRNA when autophagy was inhibited but, instead, recovered rRNA when the UPS, a more selective degradation pathway, was impeded by proteasome inhibitors (**Figure 2**). Despite rescuing rRNA levels with proteasome inhibitors, we were unsuccessful in recovering rRNA levels when the E1 ubiquitin-activating enzyme was inhibited in contradiction with the common mechanism of the UPS. One possible explanation for this observation is that RPs are being modified with a different signal molecule that then targets the ribosome for proteasome-dependent degradation. It has previously been shown through proteome-wide mapping that following 8 hr treatment with CHX that the substrate recognition subunits of a major class of E3 ubiquitin ligases known as RING E3 ubiquitin ligases are short lived (55). As such, after 24 hr treatment with CHX, most RING E3 ubiquitin ligases are presumably nonfunctional. This ineffectiveness may be more widespread than previously appreciated based on our observation that total ubiquitin protein levels are also depleted after prolonged treatment with CHX (**Figure 6**). Consequently, the ability of all E3 ubiquitin ligases to modify target proteins with ubiquitin may be further hindered after periods of sustained global translation inhibition.

Logically, given that ribosomes are being globally impacted, it stands to reason that the cell would mechanistically require a response that did not depend on the RQC pathway as many of these factors have been shown to sub-stoichiometric to ribosomes (56). One such pathway that does recognize globally stalled ribosomes is the RSR pathway. We conclude that at these extended periods of translational arrest that the RSR pathway does not influence the ribosome degradation as ribosome turnover was still observed when ZAKα protein levels were depleted or when ZAKα activity was inhibited (**Figure 9**). Although ZAKα protein levels were sufficiently reduced after siRNA knockdown, we were unable to completely abolish downstream activation of p38. Therefore, it remains possible that stimulation of an unknown upstream factor in response to ribosome stalling or activation of another factor downstream of p38 phosphorylation could be responsible for the observed ribosome turnover.

During this study, we also observed other instances unrelated to global inhibition of translation that resulted in uncategorized large-scale ribosome turnover. When A549 cells were treated for 8 hrs with LLOMe, we observed an ∼2-fold reduction in 28S rRNA and 18S rRNA levels (**Figure 11**). We hypothesized that under these conditions, ribosomes were being degraded by lysosome encapsulated RNases and cathepsins leaking out into the cytosol. Although we were able to confirm that LMP was occurring after treatment with LLOMe (**Figure 11**), we were unsuccessful in sufficiently reducing the endoribonuclease, ribonuclease T2 (RNase T2), protein levels by siRNA knockdown (data not shown) due to its low abundance in A549 cells (as indicated by The Human Protein Atlas) (57,58). As such, we were unable to confirm if RNase T2 was the nuclease responsible for the observed rRNA degradation following LMP. It should be noted that lysosomes also contain 5’-3’ exonucleases so investigation of these factors may be warranted in future studies. Additionally, we observed that A549 cells treated with TAK243 alone for 24 hrs resulted in ∼50% reduction of 28S rRNA and 18S rRNA levels (data not shown). In contrast with CHX treatment, when cells were co-treated for 24 hrs with TAK243 and MG132, rRNA levels remained depleted (data not shown). Despite ubiquitin being closely associated with the UPS, it has also been shown to be important for the regulation of autophagy (59). Therefore, it is possible that the depletion of ribosomes following TAK243 treatment is from degradation by the autophagy pathway, and this is currently being investigated.

CHX is a cytosolic inhibitor of translation and does not target mitochondrial ribosomes (60). A recent study showed that HeLa cells tolerated CHX for 1-2 days if treated during interphase but became sensitive to CHX treatment during mitosis (23). It was determined that this observation was due to changes in translation stringency between interphase and mitosis. During mitosis when the nuclear envelope is degraded, eIF1 is released from the nucleus and leads to increased start codon selection stringency, which consequently shifts the global proteome. Of the transcripts that were highly translated during mitosis, nuclear-encoded mitochondrial mRNAs were enriched. These transcripts are translated by cytosolic ribosomes and their translation was dependent on the release of nuclear eIF1. However, when nuclear eIF1 was depleted, this resulted in loss of mitochondrial membrane potential which correlates with reduction of mitochondrial fitness (23). We also observed by staining with Hoechst 33342 that the small fraction of A549 cells that underwent cell death after 24 hrs of CHX treatment had fragmented DNA (data not shown). This is indicative of endonuclease G and apoptosis-inducing factor (AIF) release from the mitochondria due to mitochondrial fragmentation (61,62). Another report using acetoxycycloheximide (E-73), a structural derivative of CHX, in human leukemia Jurkat T cells showed that treatment resulted in activation of the JNK pathway and subsequent release of cytochrome c from the mitochondria (63). Consequently, this ultimately led to caspase-dependent apoptosis. In comparison with CHX, E-73 inhibits translation 10-fold more potently, and, as such, was also able to induce apoptosis at concentrations ∼100-fold lower. It was also shown that Jurkat T cells treated with 50 μM CHX (almost 6 times lower than the concentration of CHX used in our studies) for 2 hrs had accumulation of cytochrome c in the cytosol (63). Therefore, we are currently testing to determine if prolonged treatments with high dosages of CHX result in loss of mitochondrial fitness.

## ACKNOWLEDGMENTS

Experiments were conceived and performed by PJR, CAC, and MGK with input and help from MA. PJR was supported by NIH grants T32GM086252 and T32GM141955. This work was supported by NIH grant R35GM146924 to MGK.

